# Myelinating Schwann cells and Netrin-1 control intra-nervous vascularization of the developing mouse sciatic nerve

**DOI:** 10.1101/2020.11.12.379792

**Authors:** Sonia Taïb, Noël Lamandé, Sabrina Martin, Piotr Topilko, Isabelle Brunet

## Abstract

Peripheral nerves are vascularized by a dense network of blood vessels to guarantee their complex function. Despite the crucial role of vascularization to ensure nerve homeostasis and regeneration, the mechanisms governing nerve invasion by blood vessels remain poorly understood. We found that the sciatic nerve invasion by blood vessels begins around embryonic day 16 and continues until birth. Interestingly, intra-nervous blood vessel density significantly decreases during post-natal period, starting from P10. We show that, while the axon guidance molecule Netrin-1 promotes nerve invasion by blood vessels during embryogenesis, myelinated Schwann cells negatively control intra-nervous vascularization during postnatal period.

## Introduction

Formation of vascular plexus and its maturation into a dense network of blood vessels is one of the most important mechanisms during embryonic and post-natal development in vertebrates (Kolte, McClung, and Aronow 2015). Among numerous physiological tasks, the vascular network transports oxygen and nutrients and eliminates waste products, all critical for organ survival and cellular homeostasis. During early stages of embryogenesis, by a process called vasculogenesis, a part of the mesoderm differentiates into endothelial cells, forms a lumen and deposits a basal lamina to create a vascular plexus *de novo*. To become functional, blood vessels mature and specialize by recruitment of mural cells and form a network of arteries, capillaries and veins. During development and in the adult, new blood vessels are formed from pre-existing ones by a process called angiogenesis. This mechanism involves specialized endothelial cells named tip cells. Thanks to the expression of specific receptors, tip cells can sense the microenvironment, and respond to guidance cues to lead the angiogenic sprout. Thus, angiogenesis not only allows the rapid vascularization of developing tissues and organs but ensures appropriate vascularization rate adapted to specific needs in nutrients and oxygen (Carmeliet and Jain 2011).

Peripheral nerves, connecting the central nervous system to the rest of the body, are composed of axons covered by myelinating and non-myelinating Schwann cells (SC). During neural development, axons are guided through embryonic tissues to reach their final targets (Stoeckli 2018), supported by secreted trophic factors ensuring axonal survival until the connection is ultimately stabilized when the appropriate target is reached (Ye et al. 2019). Simultaneously, SC precursors derive from the neural crest cells and migrate from the neural tube around embryonic day E10,5 to contact axons and differentiate into immature SC around E15/E16 (Woodhoo and Sommer 2008). Finally, around birth they differentiate into either myelinating or non-myelinating SC. Schwann cells are necessary to ensure axonal survival during development and also in regeneration (Jessen, Mirsky, and Lloyd 2016). Myelin sheaths are produced by SC wrapping larger axons in a 1:1 ratio to allow rapid saltatory conduction of action potentials. In rodents, myelination is progressive, starts around birth and lasts during about 2 weeks (Woodhoo and Sommer 2008). Non-myelinating SC associate with multiple small caliber axons to form Remak bundles without forming compact myelin sheaths. Adult nerves are stable structures, with nerve fibers surrounded and protected by three layers of connective tissues: the endoneurium, perineurium and epineurium (Kaplan et al. 2009). While nerves acquire a complex, multi-cellular composition, especially with connective tissue, specific metabolic needs are covered by an adapted and specific vascularization. Indeed, a dense network called the *vasa nervorum*, ensures the maintenance of proper nerve homeostasis. This is of particular importance when nerves are formed from axons covering a long distance from their soma to the innervated target. The organization of this vascular system has been previously described (Boissaud-Cooke, Pidgeon, and Tunstall 2015). In fact, blood vessels are found within the 3 connective tissues, all along the nerve length, forming the peri- and intra-nervous vascular system. They are composed of a single layer of endothelial cells surrounded by mural cells: the smooth muscle cells and pericytes allowing the contraction of the blood vessel to adapt the blood flow.

Whereas the different developmental stages of the axons, connective tissues and SC have been documented, it remains unclear when and how the *vasa nervorum* develops. A better understanding of the normal development of the peripheral nerves and their components, especially the *vasa nervorum*, is of crucial importance as nerves control many organs and tissues. Moreover, understanding the molecular control of nerve vascularization and re-vascularization after injury is of primary importance in regenerative medicine. In fact, after transection, peripheral nerves can repair and reconnect and this process implicates migration of SC to guide regrowing axons. It has been shown that blood vessels inside the nerve are used by SC to direct this migration (A.-L. Cattin et al. 2015). Moreover, nerve grafts appears to be more efficient when their vascularization is preserved, allowing a better regeneration (D′Arpa et al. 2015).

In this study, we performed a time course analysis of the intra nervous vascularization and discovered mechanisms for the development and maturation of the intra-nervous vascular system in the sciatic nerve. Importantly, we decipher the cellular and molecular actors governing the *vasa nervorum* formation.

## Results

### The intra-nervous vascular system (INV) develops rapidly from embryonic day E16

To define when the first blood vessels penetrate the nerves and mature into *vasa nervorum*, sciatic nerves were dissected and analyzed for the presence of blood vessels at successive stages starting from embryonic day 15 (E15). At this stage arteries and nerves start to be aligned in the skin (Mukouyama et al. 2002). Endothelial cells were visualized using CD31 marker in order to assess the level of nerve vascularization. At E15.5, the extrinsic artery of the primitive sciatic nerve is aligned with axons (TUJ-1 staining). However, no blood vessels were observed inside or surrounding the nerve (**Figure 1A**).

**Figure 1.**
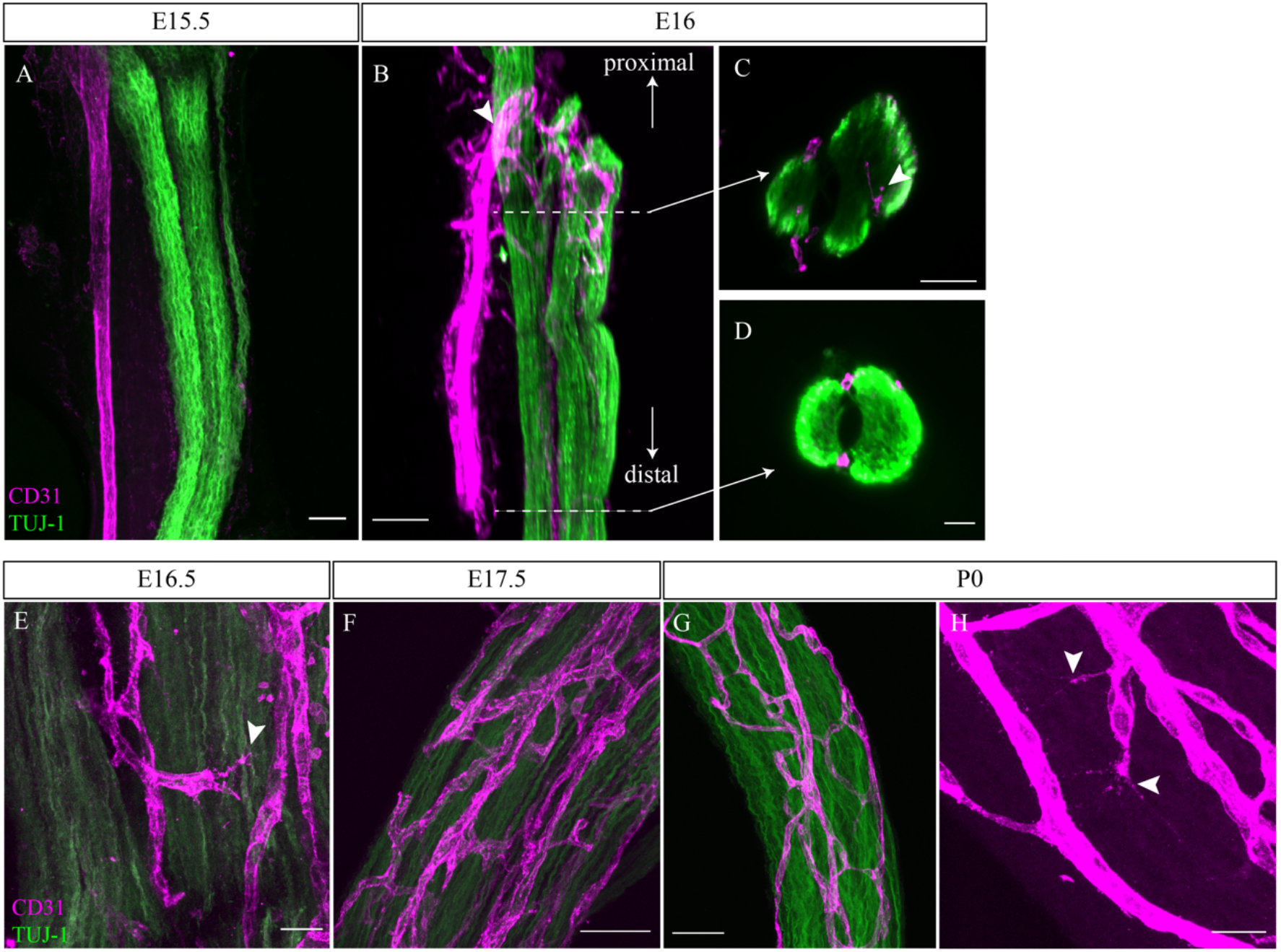
INV develops rapidly starting from embryonic day E16, angiogenesis is still ongoing at P0. **A.** Whole-mount preparation of an immature sciatic nerve (axons: TUJ-1, green) and blood vessels (endothelial cells: CD31, magenta) at embryonic day E15.5. **B.** 3D view of a primitive sciatic nerve at E16. New blood vessels (CD31, magenta) emerging from the aligned artery (arrowhead) forming the peri-nervous vascular system. **C.** Orthogonal view of the proximal part of the nerve showing blood vessels starting to form the intra-nervous vasculature (arrowhead) and of the distal part, **D**, showing no intra-nervous blood vessels. **E.** An endothelial tip cell inside a nerve from E16.5 embryo (arrowhead). **F.** At E17.5, the INV is denser. **G.** Vascularization at P0, **H** angiogenesis is still ongoing, as endothelial tip cells displaying filopodias are visible (arrowheads). Scale bars are 50 μm for A, B, C, D, F and 20 μm for E, G and H.

At E16, we noticed new blood vessels emerging from the aligned extrinsic artery to form a peri-nervous vascular plexus (**Figure 1B***, arrowhead*). Developing from this latter, numerous angiogenic sprouts invade the inside part of the immature nerve, starting to form the intra nervous vascular network (INV) as observed in the optical section of the proximal part of the nerve (**Figure 1C***, arrowhead*). However, at E16, in the distal part no blood vessels were found inside the nerve (**Figure 1D**), suggesting a proximo-distal gradient of nerve invasion by blood vessels. At E16.5, the INV continues to form new blood vessels by angiogenesis, with endothelial tip cells formation (**Figure 1E***, arrowhead*) leading to a denser, anastomotic vascular network at E17.5 (**Figure 1F**). Finally, at birth (P0), very few tip-cells were found inside the nerve (**Figure 1H**, *arrowheads*), suggesting that sciatic nerve vascularization takes place before birth (**Figure 1G and H**).

### Blood vessels inside the nerve mature during post-natal development, undergoing arterial differentiation and pericytes recruitment

To be fully functional, blood vessels have to be mature and organized as a hierarchical vascular network. Endothelial cells of arterioles specifically express the protein connexin-40 (Cx-40) and are covered by smooth muscles cells (SMC) to control their diameter and adapt the blood flow. The extrinsic artery, expressing Cx-40, visualized by GFP expression, and covered by SMC, in magenta, (**Figure 2A***, arrowhead*) is divided into continuous ascending and descending arterioles to directly supply each region of the nerve and ensure arterial blood flow to the entire nerve. Pre-existing blood vessels of the INV also undergo arterial differentiation, as observed in sciatic nerve at P5. Endothelial cells expressing Cx-40 in an isolated manner were visible (**Figure 2B***, arrowheads*), suggesting that new arterial branches undergo differentiation inside the nerve. Notably, arterialization of the penetrating vessels is still ongoing as some parts are already covered by SMC whereas newly differentiated arteries have still not recruited SMC (**Figure 2A**). Pericytes, expressing the protein NG2 and covering endothelial cells, are important for maintenance of the blood-nerve barrier and are able to contract to change the capillary diameter (Bergers and Song 2005). At P0, numerous endothelial cells of the INV are already covered by pericytes (**Figure 2C**). Some of these cells cover new blood vessel branching (**Figure 2D**) as it has been already reported that pericytes are critical for angiogenesis (Bergers and Song 2005). These pericytes at P0 have an elongated shape and their coverage is relatively sparse compared to the pericytes observed at P10 (**Figure 2E**). Thus, at birth, INV already displays vessel wall maturation that will remain during nerve postnatal morphogenesis.

**Figure 2.**
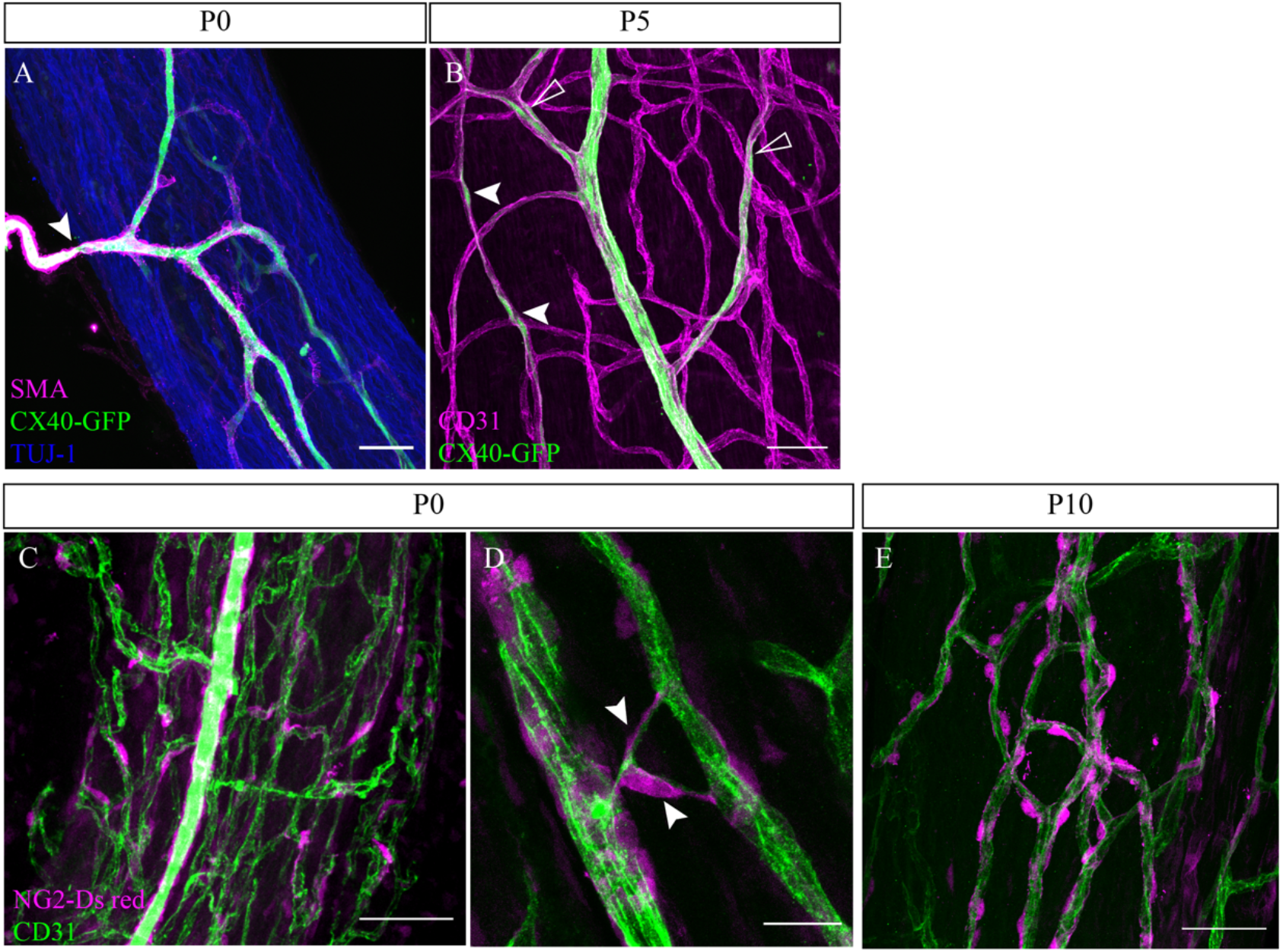
INV matures during post-natal stages with arterial differentiation and pericytes recruitment. **A.** Whole-mount staining of a P0 sciatic nerve from a Cx40-GFP reporter mouse showing an artery composed of endothelial cells expressing the protein CONNEXIN-40 (in green) and covered by smooth muscle cells (stained in magenta, smooth muscle actin staining (SMA)). This artery has invaded the sciatic nerve (axons expressing TUJ-1, blue). **B.** At P5, some endothelial cells express the protein connexin-40 (white arrowheads) and new arterial branches are differentiating (empty arrowheads). **C.** NG2-Dsred mice were used to observe pericytes (red) covering endothelial cells (CD31, green) of a P0 sciatic nerve. **D.** Close-up view of a P0 sciatic nerve vasculature showing pericytes covering new blood vessel branching (arrowheads). **E.** Pericytes coverage of the INV at P10. Scale bars are 50 μm for A, B, C, E and 20 μm for D.

### INV density decreases during post-natal development

Post-natal maturation of the peripheral nerves includes the production of the myelin sheaths by SC. Myelination starts at birth, continues for about 2-3 weeks and increases both nerve size and caliber (Jessen and Mirsky 2005).

We hypothesized that the growth of the nerve would provoke an increase of the INV during post-natal development to adapt the nerve supply in oxygen and nutrients. To test this hypothesis, we analyzed CD31 immunoreactivity in sciatic nerves cross sections from P0 (**Figure 3A**) to young adult stage (8 weeks old mice) (**Figure 3B**).

**Figure 3.**
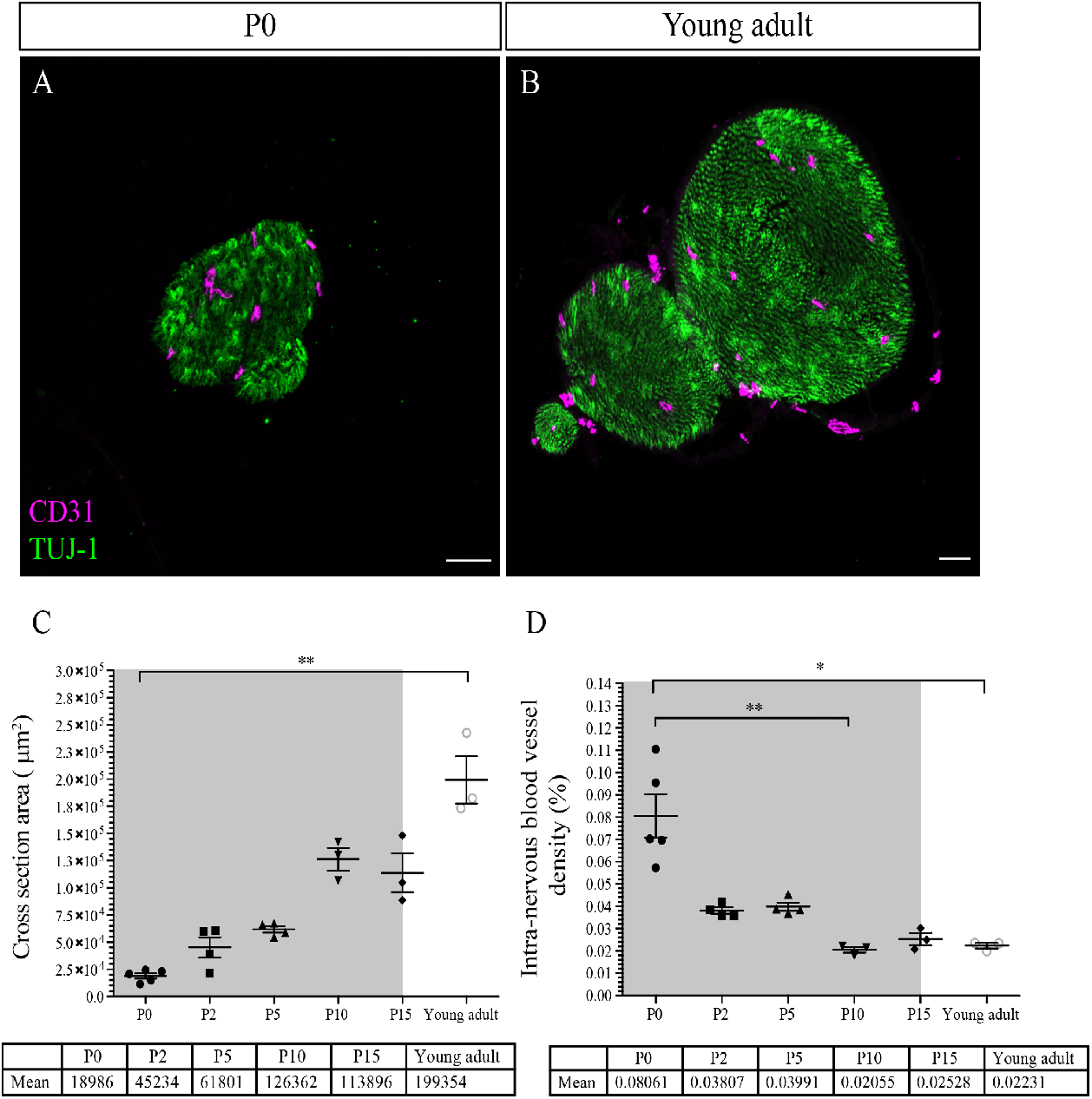
INV density decreases during post-natal development. **A.** Sciatic nerve’s transversal section at P0 from a young adult mouse (**B.**) showing blood vessels (CD31, magenta) and axons (TUJ-1, green) inside the nerve in and **C.** Quantification of the surface area of nerves’ cross section, from P0 to young adult stage (around P54). The grey box represents the myelination period from P0 to P15. **D.** Intra-nervous CD31 density expressed as percentage of total nerve cross section area. Only the blood vessels inside the nerve were counted. Blood vessel’s density decreases significantly from P0 to P10 and stabilizes. (n=3-5 animals for each stage, >50 cross sections were analyzed per nerve, mean ± SEM, Kruskal-Wallis and Dunn’s multiple comparisons test, *p<0,05, **p<0,01).

As expected, the caliber of the sciatic nerve gradually increases during post-natal development by a factor of 10 from P0 to young adulthood (**Figure 3C**). Area densities of CD31 immunoreactivity was quantified as percentage of total nerve area and appears to vary overtime (**Figure 3D**). The intra-nervous CD31 positive area covers around 8% of the nerve at P0. this value decreases progressively after birth and stabilizes after P10, to reach around 2% of nerve coverage, and does not change later at young adult stage. Interestingly, this decrease correlates with myelination period during the 2 weeks after birth (grey box, **Figure 3D**).

### Ablation of myelinating Schwann cells leads to an abnormal vascularization of the sciatic nerve

As nerve vascularization density decreases during post-natal development and while myelin is produced and since the majority of cells within the sciatic nerve are SC (∼70%) (Stierli et al. 2018), we wondered if myelinating SC could affect nerve vascularization. It has been previously described that SC promote endothelial cells migration *in vitro* (Ramos et al. 2015). To address *in vivo* the question of whether these cells have a role in the development of the INV, we used a transgenic mouse line in which genetic ablation of SC precursors was provoked. The transcription factor KROX20, expressed by immature SC, represents the master regulator of myelin genes and thus controls myelination formation and maintenance (Topilko, Schneider-Maunoury, Levi, Baron-Van Evercooren, et al. 1994). We crossed *Wnt1-Cre*/+ mice (Joseph et al. 2004) with *Krox20*^*GFP(DT)*/+^ mice (Vermeren et al. 2003) to obtain *Wnt1-Cre;Krox20*^*GFP(DT)*/+^ pups in which the A chain of the Diphteria Toxin (DT) is only expressed upon cre-mediated recombination, to lead to a specific death of immature SC expressing Krox20 around E15/E16. In fact, at P0 SC (normally expressing SOX10) are absent in sciatic nerves from *Wnt1-Cre;Krox20*^*GFP(DT)*/+^ mutant mice (**Figure S1 A and A’**) and at P5 myelinating SC (normally expressing S100β protein) are lacking (**Figure S1 B and B’**). These animals survive only few days after birth. Therefore, we analyzed the level of vascularization of sciatic nerves at P5. As expected, the sciatic nerves of *Wnt1-Cre;Krox20*^*GFP(DT)*/+^ mice were thinner as compared to *Wnt1-Cre* control littermates (**Figure 4A and B**). This was due to the lack of SC and their myelin sheaths, which normally considerably increase the nerve’s caliber. Furthermore, the INV network is disrupted, denser and more anastomotic, resembling embryonic immature plexus (**Figure 4C**). To further analyze the consequences of the SC ablation on the INV, we performed 3D reconstruction of this vasculature *in toto* (**Figure 4D and E**). We quantified the total length of the vasculature tree in blue and the number of blood vessel branching points in red. Those values were normalized to the nerve area. Sciatic nerves of mutant pups display an increased total length of blood vessel network (**Figure 4F**) with more branches (**Figure 4G and H**) compared to the control littermates. Altogether, these results suggest that myelinating SC could control INV development and maturation probably by inhibiting angiogenesis or related processes. Nonetheless, this phenotype could be caused by the lack of SC or by the lack of myelin.

**Figure 4.**
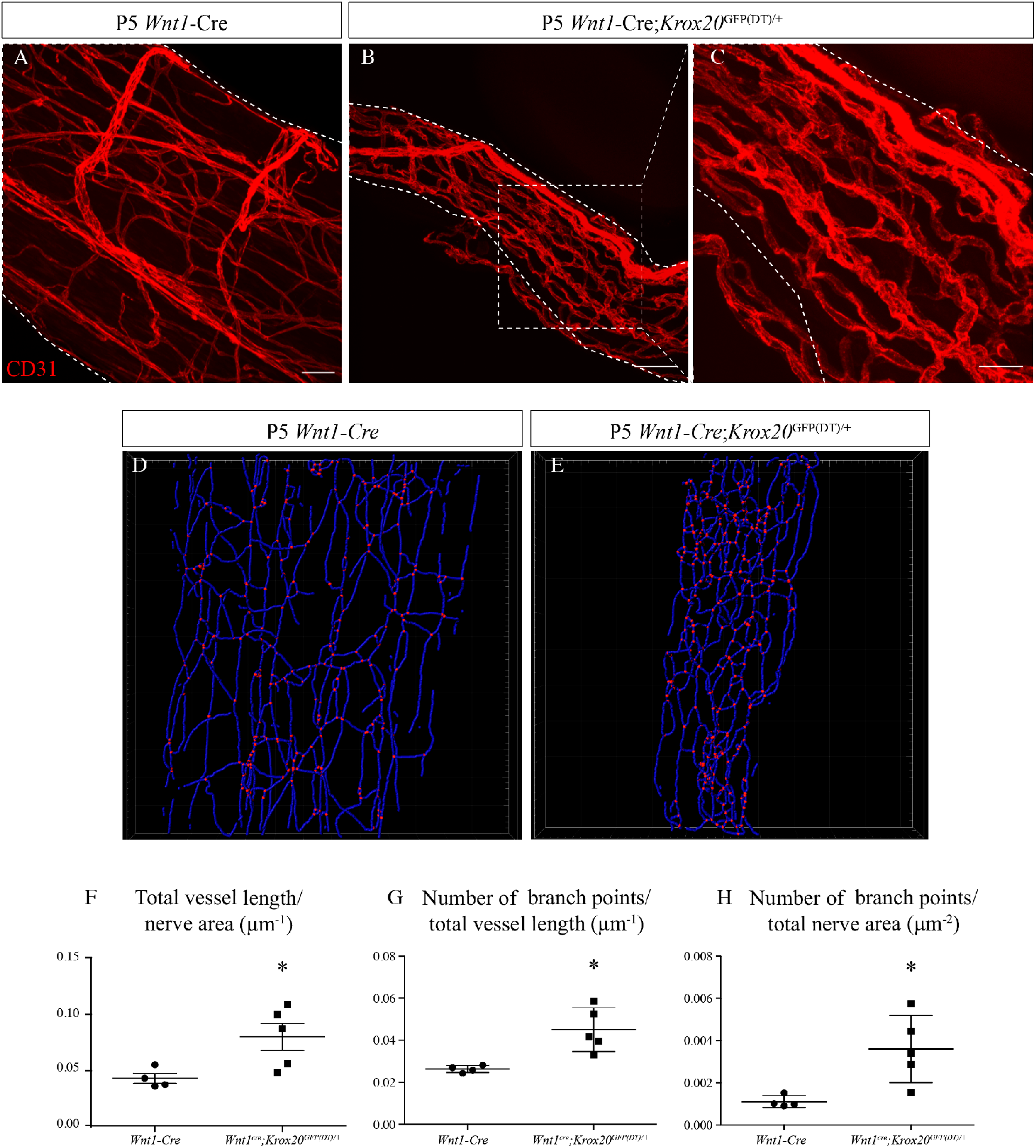
Schwann cells ablation disrupt sciatic nerve vasculature. **A.** Whole mount staining of a sciatic nerve from the control *Wnt1*-Cre mouse at P5, the blood vessels are stained with CD31 and shown in red. **B.** Vasculature of a sciatic nerve from a *Wnt1*-Cre;*Krox20*^GFP(DT)/+^ mouse at P5. **C.** Close-view showing the disruption of the blood vessels’ organization. (**D and E**) *Imaris* 3D reconstruction of sciatic nerve vasculature in blue, the branching points are shown in red. **F.** Quantification of total blood vessel length/ nerve area. **G.** Number of branching points per total vessel length. **H.** Number of branching points per nerve area. (n= per group, graphs show mean ± SEM, Mann-Whitney test, *p<0,05). Scale bars are 50 μm for A and B and 20 μm for C.

### Inhibition of myelin production provokes hypervascularization of the sciatic nerve

To decipher the role of the myelin sheath, independently of SC, we used the *Krox20*^*Cre/Fl*^ mouse line. As previously described, in these mice, *Krox20* from the floxed allele is expressed until enough Cre recombinase accumulates resulting in the delayed inactivation of the *Krox20* gene in myelinating SC (Decker et al. 2006). Thus, *Krox20* inactivation blocks SC at an early stage of their differentiation and therefore prevents the formation of myelin sheaths (**Figure 5A**). *Krox20*^*cre/Fl*^ mutants survive around 20 days after birth and show tremors. We chose to analyze sciatic nerves before lethality, at P19, so that the sciatic nerves from control pups are fully myelinated. The vasculature of sciatic nerves from control and mutant mice were compared using CD31 staining on whole mount preparations. Interestingly, sciatic nerves of mutant mice appears to be more densely vascularized compared to control littermates (**Figure 5B and B’**). Thus, we performed 3D reconstruction of the INV (**Figure 5C and C’**). As compared to control mice, quantifications revealed a higher blood vessel total length (**Figure 5D**) and higher number of branching points in mutant’s sciatic nerves (**Figure 5E and F**). We then aimed to better characterize vascular network defects and assessed maturation and permeability of the INV. Endothelial cells composing the INV express claudin-5, a tight junction protein, important for the acquisition of a barrier property (Peltonen, Alanne, and Peltonen 2013). At P18, in sciatic nerves from control mice, Claudin-5 staining colocalized with CD31 staining in almost all blood vessels (**Figure 5G**). In sciatic nerves from *Krox20*^*Cre/Fl*^ mice, Claudin-5 is poorly expressed by endothelial cells as almost all blood vessels branches are Claudin-5 negative (**Figure 5G’**, *arrowheads*).

**Figure 5.**
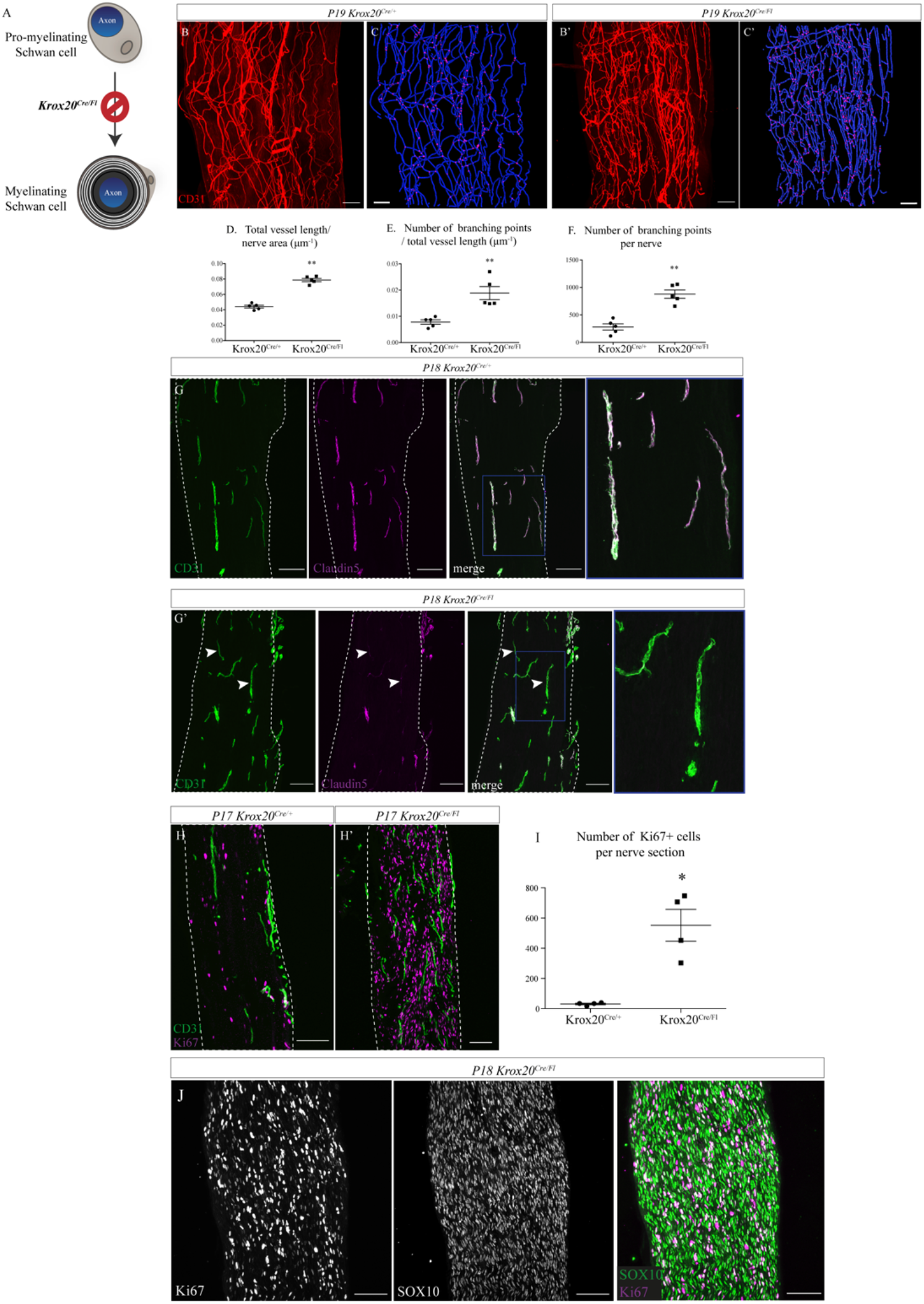
Sciatic nerve is hypervascularized upon myelination inhibition. **A.** Schematic illustrating *Krox20* knock out leading to myelination inhibition. **B and B’**. Whole mount staining of sciatic nerves from a *Krox20*^*cre*/+^ control mouse and from a *Krox20*^*cre/Fl*^ mouse at P19, blood vessels are in red. **C and C’.** Imaris 3D reconstruction of sciatic nerve vasculature in blue, the branching points are in pink. **D.** Quantification of total blood vessel length/ nerve area. **E.** Number of branching points over total vessel length. **F.** Number of branching points per nerve. (n=5 per group, graphs show mean ± SEM, Mann-Whitney test, **p<0,01). **G and G’.** Longitudinal sections of sciatic nerve dissected *Krox20*^*cre*/+^ and *Krox20*^*cre/Fl*^, at P18. Endothelial cells expressing CD31 are in green and claudin5 in magenta. Dotted lines represent the borders of the nerve section. Some vessel branches do not express claudin5 (arrowheads) (**H and H’**) Longitudinal sections of sciatic nerves dissected from a *Krox20*^*cre*/+^ mouse and *Krox20*^*cre/Fl*^, at P17. Cells in proliferation express the protein Ki67 (magenta) and blood vessels are in green. **I.** Quantification of Ki67 positive cells per nerve section. n=4 animals per group, around 10 sections were quantified per nerve, graph shows means ± SEM, Mann-Whitney test, *p<0,05. **J.** SOX10 and Ki67 staining on sciatic nerve longitudinal sections. Scale bars are 100 μm.

Altogether, those data suggest that myelin sheath controls sciatic nerve vascularization by inhibiting angiogenesis and by limiting vascular rate and maturation.

We checked, using Ki67 staining, if the endothelial cells were proliferating and still producing new blood vessels. Ki67+ cells were not endothelial cells, proof that blood vessels have stopped proliferating. This finding corroborate our previous data about vascularization rate at P15/young adult (**Figure 2**). Interestingly, the number of proliferating cells is considerably higher in the unmyelinated sciatic nerves (**Figure 5H, H’ and I**). As reported before (Topilko, Schneider-Maunoury, Levi, Baron-Van Evercooren, Chennoufi, et al. 1994), these proliferating cells are SC, as they specifically express the protein SOX10 (**Figure 5J**). Indeed, activation of the genes controlling myelination, such as KROX20, induces the arrest of the proliferation state of SC in favor of differentiation and production of myelin sheath (Salzer 2015).

### The guidance molecule Netrin-1 is involved in INV formation

We next investigated the molecular control of INV development at embryonic stages. Netrin-1 is a guidance molecule known to control axonal guidance and also angiogenesis (Park et al. 2004). Studies have shown that it can also be implicated in nerve regeneration, stimulating axonal and blood vessels regrowth (Dun and Parkinson 2017), (Madison, Zomorodi, and Robinson 2000). Therefore, we asked whether Netrin-1 could regulate sciatic nerve vascularization during normal development. As we described, vascularization of sciatic nerves is still ongoing around birth, involving active angiogenesis. First, we used *Ntn1*^*lacZ*/+^ knock-in mice to report Netrin-1 protein expression in the nerve. At P2, β-galactosidase activity, reporting Netrin-1 expression, was found close to blood vessels, in areas compatible with a potential role of Netrin-1 in endothelial cell guidance (**Figure 6A**). As *Ntn1*^−/−^ mice die around birth (Serafini et al. 1996), we analyzed *Ntn1*^*lacZ/lacZ*^ knock-in embryos and explored sciatic nerves vascularization of embryos at E17.5, when INV has already initiated. Embryonic sciatic nerves were dissected and we assessed blood vessels quantity using CD31 staining (**Figure 6B and B’**). Whereas there is no difference in nerve size (**Figure 6C**), we found that nerves from mutant embryos are less vascularized as compared to control embryos (**Figure 6D**). To confirm this result, and to make sure that the dissection did not alter the nerve integrity and vascularization, we quantified nerve vasculature at this stage directly within the embryonic limb. Embryos’ limbs were entirely dissected, as they contain the sciatic nerve (**Figure 6E and 6E’**, left top corner, dotted area). We performed whole-mount immunostaining of the entire limb to stain axons (neurofilament) and blood vessels (CD31). Limbs were then cleared using an adapted iDISCO+ protocol (Renier et al. 2014) and the areas of interest were imaged in 3- dimensions. Using *Imaris* software, we observed the sciatic nerve inside the limb, thus keeping the physical integrity of the nerve and its surrounding (**Figure 6F and 6F’**). We also found that, whereas sciatic nerves of *Ntn1*^*lacZ/lacZ*^ embryos had the same size as control littermates (**Figure 6G**), they were less vascularized. This was confirmed by quantification of CD31 positive area density inside the nerve (**Figure 6H**), with a similar ratio and in agreement with our findings on dissected nerves. This data suggests that Netrin-1 is likely to be one of the guidance molecules responsible for the attraction and angiogenesis of the blood vessels inside the embryonic nerve.

**Fig. 6.**
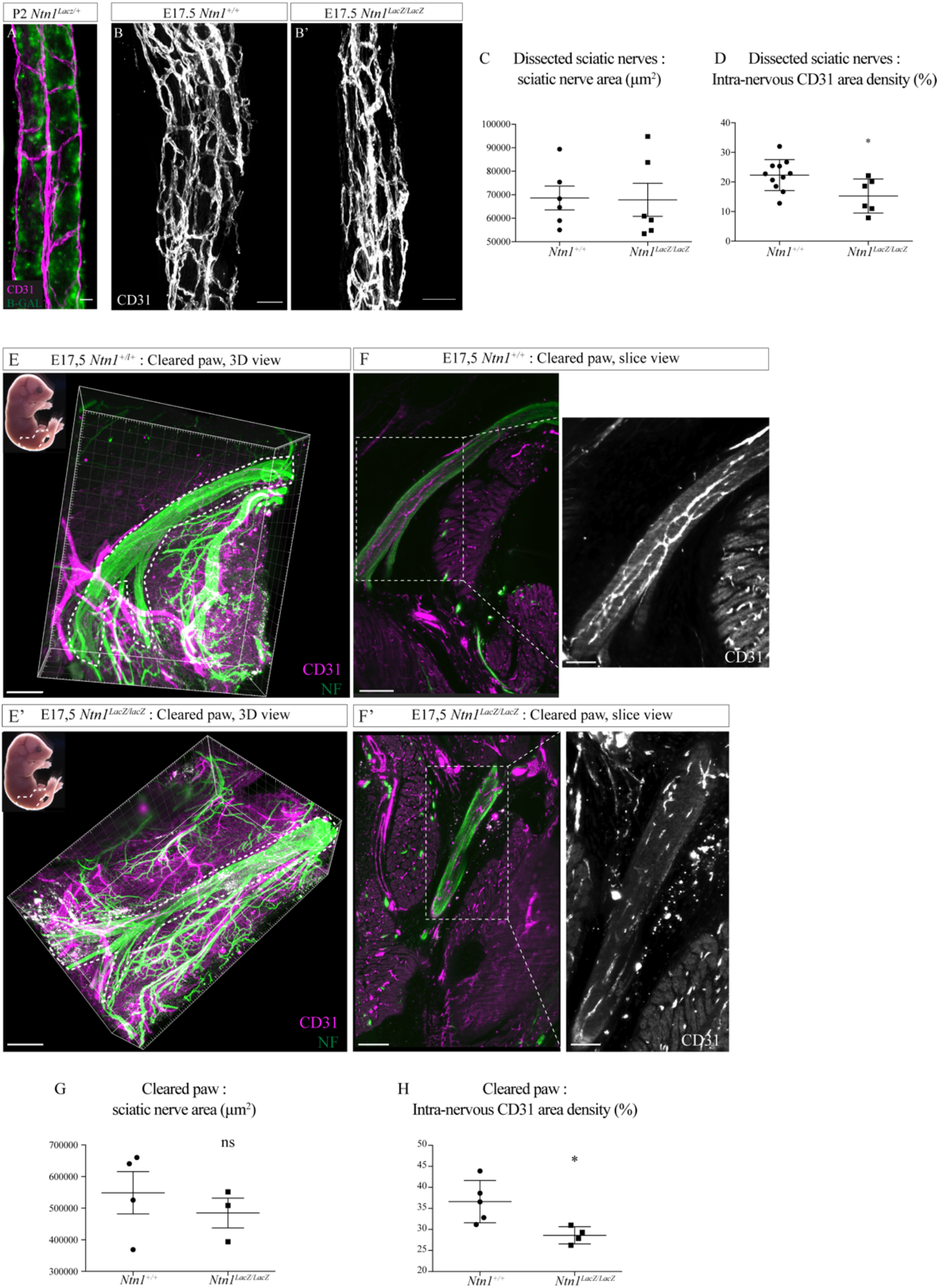
Netrin-1 hypomorph embryos display a reduced intra-nervous vascularization. **A.** X-Gal staining (green) of a nerve whole-mount from a P2 *Ntn1*^*lacZ*/+^ mice and vasculature in magenta (CD31). **B.** CD31 whole-mount staining of a sciatic nerve from E17.5 *Ntn1*^+/+^ and *Ntn1*^*LacZ/LacZ*^ embryos (maximum intensity projection). **C.** Quantification of nerve area (from NF staining). **D.** Quantification of the intra-nervous CD31-positive total area after images thresholding, showing that vascularization is significantly reduced in mutant embryos. **E and E’.** Top left corner: E17.5 embryos (the dotted line represents the dissected limb containing the primitive sciatic nerve) and 3D view from *Imaris* software of the embryo’s limb, after CD31 (magenta) and neurofilament (NF, green) staining and clearing protocol (iDISCO+). Dotted lines delimit the immature sciatic nerve. **F and F’.** Slice view of the cleared limb allowing the observation of the blood vessels inside the sciatic nerve. **G.** Quantification of nerve area (from NF staining)). **H.** Quantification of intra-nervous CD31-positive area, using optical slices from cleared E17.5 embryos’ limbs. n=4-5, Mean ± SEM, Mann-Whitney test, *p<0,05. Scale bar is 20 μm for A, 50 μm for B, 80 μm for D and D’, 200 μm for E and E’.

## Discussion

In the developing peripheral nerves, blood vessel formation is tightly controlled, both spatially and temporally, as the INV has to be well-adapted to maintain homeostasis and ensure proper function. This study provides the time-course of the development and maturation of the mouse sciatic nerve’s INV. We find that INV appears relatively late during embryonic development, around E16, as compared to the vasculature of the central nervous system (CNS) which begins around E9,5 (Tata, Ruhrberg, and Fantin 2015) (Himmels et al. 2017) to ensure adequate delivery of oxygen and nutrients to neural progenitors. This temporal difference lies in the specific timing of CNS morphogenesis that starts earlier and maturates over a longer time. E16 in the mouse embryo represents a turning point for the peripheral nervous system (PNS) as SC precursors acquires an immature phenotype (Jessen and Mirsky 2019). Interestingly, this step matches the apparition of the first blood vessels inside the nerve, suggesting there may be different molecular cues, expressed by immature SC participating in the attraction of blood vessels into the nerve.

Microvascular density is adapted to tissue oxygen and metabolic needs. During embryonic and post-natal development, nerves gain size and volume, due to myelination and production of the connective tissues. We hypothesized that this leads to increased blood vessel density, mainly based on potential formation of hypoxic zones inside the nerve during nerve growth driving angiogenesis. Interestingly, we found the opposite: the vascular tree stabilizes, angiogenesis is stopped, or remains very low, and as a consequence INV density decreases during post-natal development. This suggests that nerves reach their optimal blood vessel density around P10 similarly to the CNS in which angiogenesis continues until P10 in mice (Harb et al. 2013). At P5, there is still angiogenesis ongoing but at a lower rate. Thus, myelination rate is faster than vascularization, leading to vascular density decrease and stabilization around P10, when both process seem to end.

Even though SC and blood vessels are significant players in peripheral neuropathies pathogenesis and nerve repair mechanism (A. L. Cattin et al. 2015), no studies are available regarding their relationship during physiologic development. In the developing PNS, the vascularization needs to be tightly controlled and we have shown that SC have a central role. To our knowledge, we here provide the first evidence that myelinating SC control the vascularization of the sciatic nerve during development. As the major phenomenon of the post-natal development of the PNS is myelination and that concomitantly the INV density is decreasing, we hypothesized that myelinating SC may be responsible for the blocking of angiogenesis. We were able to discriminate specific role of SC and myelination by genetically ablating either SC or myelin production. In both cases, we found that this ablation led to a hypervascularized nerve. Nevertheless, we observed that SC death provokes a more disrupted and immature INV, as compared to the phenotype of sciatic nerves only lacking myelin sheaths. Mukouyama and colleagues have shown that SC are required for arterial differentiation in the skin of mouse embryos (Mukouyama et al., 2002). This suggests that SC bodies provide physical and/or chemical cues that control blood vessel maturation. This is also what we observe as blood vessels of the INV failed to express “barrier proteins” such as Claudin-5 when myelin production was inhibited. Another study has suggested that SC-conditioned medium inhibits endothelial cell proliferation and migration *in vitro* (Huang et al. 2000) whereas another found that SC stimulates endothelial cell migration *in vitro* (Ramos et al. 2015). Since inhibition of myelination alone and total ablation of myelinating SC both led to hypervascularized sciatic nerves, this suggest that *in vivo*, SC may have in fact an anti-angiogenic effect on endothelial cells during post-natal development possibly by producing anti-angiogenic molecules. Moreover, compact myelin sheaths are dense structures, composed of lipids (Salzer 2015), and could have biophysical properties not favorable or less permissive to the migration and propagation of blood vessels. This environment could also block the diffusion of pro-angiogenic factors produced by other cells in the nerve such as vascular endothelial growth factor (VEGF) expressed by neurons (Li et al. 2013).

Altogether, our data demonstrate that different stages of SC development orchestrate the INV formation: around E16 when the immature SC proliferate, INV starts to develop. Between P0 and P10, when myelination is happening, INV density decreases but maturation occurs. This anti-angiogenic effect of the glial cells appears to be the opposite in the CNS. Interestingly, it has been shown that, oligodendrocytes promote angiogenesis and endothelial cell proliferation in the white matter (Yuen et al. 2014).

Another question raised by our findings is do SC and myelin have a role in the maintenance of the INV? Since *Wnt1-Cre;Krox20*^*GFP(DT)*/+^ and *Krox20*^*cre/Fl*^ mice die soon after birth (around P5 and P20, respectively), we could not study further in time the role of SC and myelin regarding the maintenance of the INV. To gain insights on this and to know if myelin is not only necessary for INV onset, but also for its maintenance, futures studies could be done in mice models of demyelination at adult stage using PLPCre-ERT2; Krox20^fl/fl^ animals. This could also provide useful information on the role of INV in degenerative diseases in which myelin is targeted, as INV maintenance could perhaps delay disease progression by maintaining nerve homeostasis.

Guidance molecules controlling angiogenesis in different systems during embryogenesis are reported (Michaelis 2014) but the molecular control of INV formation is not well-known. We chose to focus on Netrin-1, a protein known to play a role in migration of different cell types, including endothelial cell, during development (Bradford, Cole, and Cooper 2009). The role of Netrin-1 regarding angiogenesis remains controversial, and seems to be dual, as it is in axon guidance, depending on the receptor it binds to (Yang et al. 2007). It has been shown that Netrin-1 stimulates angiogenesis *in vitro* and also *in vivo*, in the retina (Park et al. 2004). Despite the possible involvement of other factors, loss of Netrin-1 function was sufficient to induce defects in INV formation, even though this was not sufficient to completely block nerve’s vascularization. Indeed in our study, Netrin-1 hypomorph provokes hypovascularized sciatic nerves at E17,5, indicating that Netrin-1 is involved in INV establishment.

Since the formation of INV is finely regulated, molecularly and cellularly, by actors we have identified, the data presented here suggest a novel role for Netrin-1 and SCs during the coordinated wiring of the nervous and vascular systems. Our study opens the exciting possibility that INV and SC/myelin could be linked not only during development but also during peripheral nerve diseases progression. Thus, INV implication in demyelinating diseases should not be over looked.

---

**Figure S1.**
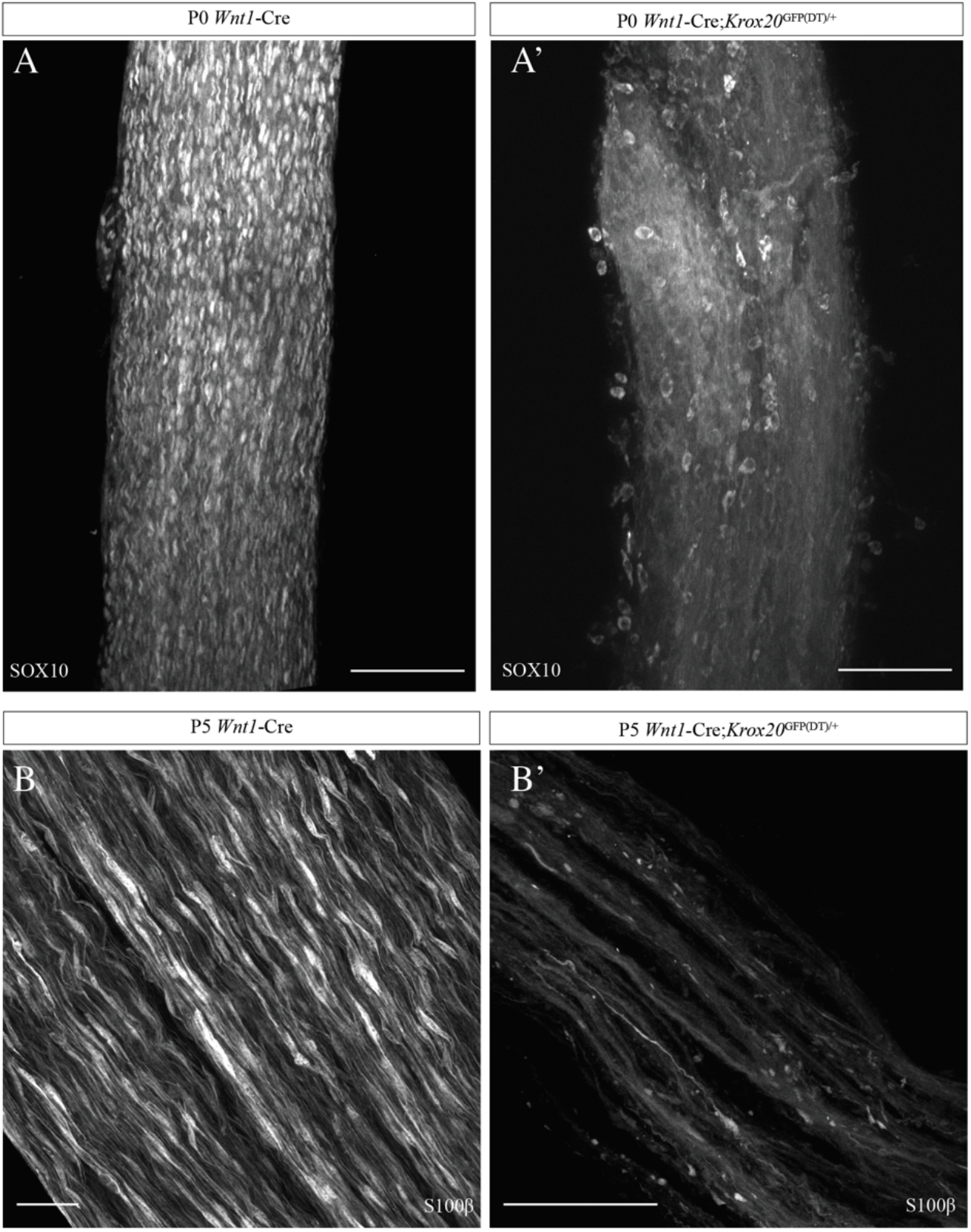
Schwann cells genetic ablation in Wnt1-Cre;Krox20GFP(DT)/+ mutant mice. **A and A’.** Whole mount immunofluorescent staining of sciatic nerves from control *Wnt1*-Cre mouse and *Wnt1*-Cre;*Krox20*^GFP(DT)/+^ mutant at P0, SC expressing SOX10 are visible in white. **B and B’.**Whole mount immunofluorescent staining of sciatic nerves from control *Wnt1*-Cre mouse and *Wnt1*-Cre;*Krox20*^GFP(DT)/+^ mutant at P5, myelinating SC expressing S100β are labelled in white. Scale bars are 100 μm for A and A’, 50 μm for B and B’.

## Materials and methods

### Animals

C57BL/6 (Janvier labs, France) mice were used for this study. *Cx40-GFP* (Miquerol et al. 2004), *Ng2-DsRed* (Tg(Cspg4-DsRed.T1)1Akik/J; Jackson laboratory) mice were previously described and genotyped with epifluorescence microscope. *Krox20*^*GFP(DT)*^ (Vermeren et al. 2003), *Krox-20-Cre* (Egr2tm2(cre)Pch/J; Jackson laboratory), *Wnt1-Cre* (129S4.Cg-E2f1Tg(Wnt1-cre)2Sor/J; Jackson laboratory), *Krox20*^*fl*/+^ (Taillebourget al. 2002) and *Ntn1^LacZ/+^*(Serafini et al. 1996) mice were previously described and genotyped by PCR. For embryonic stages, the day of the vaginal plug was counted as E0.5.

### Ethical approval

Experiments and techniques reported here complied with the ethical rules of the French agency for animal experimentation.

### Immunofluorescent staining

After euthanasia, sciatic nerves were dissected and fixed in a 4% PFA solution during 30 min at room temperature. The whole-mounts were incubated in a TNBT solution composed of Tris HCl pH 7,4/NaCl 5M/0,5% blocking reagent (Perkin)/ 0,5% Triton X-100, overnight at 4°C. Primary antibodies were diluted in the same solution, overnight at 4°C. After washes with TNT solution (Tris pH 7,4/ NaCl 5M/0,05% Triton X-100), nerves were incubated with secondary antibodies, diluted in TNBT solution, during 3 hours at room temperature. For cryostat sections: sciatic nerves, immediately after being dissected, were embedded in OCT media and snap-freezed in liquid nitrogen. Cryostat sections (14 μm) were fixed using ice-cold 100% methanol during 8 minutes then incubated in a blocking solution composed of 0,25% Triton X-100/10% fetal bovine serum/PBS during 30 minutes at room temperature. Primary antibodies were diluted in the same solution and sections were incubated overnight at 4°C. After PBS washes, secondary antibodies were diluted in a 0,1% Triton X-100/1% FBS/PBS solution and sections were incubated during 2 hours at room temperature. All the antibodies used in this study together with the information regarding their use, are listed in *Table 1*.

**Table 1.**

| Designation | Supplier and reference | Host | Dilution |
| --- | --- | --- | --- |
| CD31 | BD Pharmigen 553370 | Rat | 1:400 |
| Tuj-1 | R&D BAM1195 | Mouse<br>biotinylated | 1:500 |
| SMA-cy3 | Sigma C6198 | Mouse<br>monoclonal | 1:500 |
| Neurofilament Heavy<br>chain | Abcam ab4680 | Chicken | 1:1000 |
| Ki67 | Abcam ab15580 | Rabbit | 1:500 |
| Sox10 | Proteintech 66786-1-ig | Mouse<br>IgG2a | 1:200 |
| S100β | Proteintech 15146-1-AP | Rabbit | 1:400 |
| Goat anti-Rat IgG (H+L)<br>Alexa Fluor 555 | Thermofisher A21434 | Goat | 1:400 |
| Goat anti-Chicken IgG<br>(H+L) Alexa Fluor 647 | Thermofisher A32933 | Goat | 1:400 |
| Donkey anti-Rabbit IgG<br>(H+L) Alexa Fluor 488 | Thermofisher A21206 | Donkey | 1:400 |

### Tissue clearing

We used the iDisco + clearing method (Renier et al. 2014). Samples were first dehydrated with Methanol/H_2_O series (20%, 40%, 60%, 80% and 100%; 1 hour each) at room temperature and then incubated in 66% dichloromethane (DCM, Sigma Aldrich)/33% Methanol during 3 hours. This was followed by an incubation with 100% DCM 15 minutes, twice. Samples were then cleared with dibenzyl ether (DBE, Sigma Aldrich), overnight.

### Imaging and images processing

The images of sections were taken using Zeiss Axiozoom apotome (and associated Zen software) and Leica confocal microscope. Maximum projections of the acquired stacks were obtained with Fiji. The cleared tissues images were acquired using a light-sheet microscope and Inspector pro software (Lavision biotec). The 3D reconstruction of nerves and limbs were visualized with Imaris software (Bitplane).

### Quantifications

The density of intra-nervous blood vessels on sciatic nerve cross sections, at different ages, was quantified using the same semi-automatic method on Fiji for all the sections. After thresholding the images and creating a mask for the channel representing the blood vessels’ staining, the total area contained in the section was quantified. This value was reported to the area of the section. Quantification of the length and number of branch points of the vasculature was done using “Surface” and “Filament” tools of Imaris software.

For Ntn1^+/+^ and Ntn1^LacZ/LacZ^ embryos, quantification of the nerve area (neurofilament staining) and intra-nervous blood vessel density (CD31 staining) were made using Fiji software.

### Statistical analysis

Data shown are expressed as means ± SEM. Graphs and statistical analysis were performed using Graphpad Prism software. The different statistical tests used were specified in the figures’ legends.

## Acknowledgments

We acknowledge Philippe Mailly for the help in image analyses.

